# Sensory cortical disinhibition is required for long-term fear memory in humans

**DOI:** 10.64898/2026.08.28.745592

**Authors:** Joshua A Brown, Yijia Ma, Yifan Wang, John J Massa, Peter Kuan-Hao Cheng, Chaowen Chen, Mingzhou Ding, Norman B Schmidt, Mohammed R Milad, Jair C Soares, Raymond J Dolan, Chenyu You, Wen Li

## Abstract

The transformation of transient threats into enduring memories is fundamental to survival, yet how the human brain constructs long-term fear memory remains elusive. Early sensory cortex, traditionally viewed as a passive feature analyzer, is increasingly recognized as an active participant in fear processing. Here, using a multimodal approach combining high-definition α-frequency transcranial alternating current stimulation (HD α-tACS) with simultaneous EEG-fMRI and transformer-based neural decoding, we tracked fear memory development across early formation, consolidation, and long-term retention stages (assessed at 15 min, 24 h, and 7 days post-conditioning). Spatiotemporal memory representations were evident at all three time points, beginning at an early sensory stage (40-60 ms post-stimulus). Conditioning potentiated occipital α-oscillation desynchronization (a marker of visual cortical disinhibition), which predicted the strength of subsequent long-term memory traces. Critically, targeted α-tACS over early visual cortex induced V1/V2 inhibition, abolished conditioning-potentiated α desynchronization, and selectively disrupted fear-memory consolidation and retention while leaving initial acquisition intact. These findings establish sensory cortical disinhibition as a necessary mechanism for long-term fear memory in humans and identify early sensory cortex as a causal node for interventions targeting persistent maladaptive fear.

---

Fear learning and memory drive anxiety- and trauma-related disorders, such as posttraumatic stress disorder ^1^. Human research to date has largely focused on fear learning, identifying limbic circuits (including the salience network) as key neural substrates ^2,3^. Beyond initial learning, how a fear experience solidifies into a persistent memory, rather than fading as a momentary threat, lies at the core of anxiety-related pathology. However, the mechanistic principles governing the formation of long-term fear memory in the human brain remain unclear.

Accumulating evidence implicates the sensory cortex as a key, extra-limbic contributor to fear memory ^4–7^. Echoing early findings of sensory cortical plasticity following fear experiences in animals ^8,9^ and humans ^10–12^, a burgeoning literature has established the sensory cortex as an active participant in fear processing rather than a passive perceptual relay ^13–15^. In particular, cue-specific sensory cortex has been increasingly linked to the development and storage of enduring (days to weeks) fear memories in animals ^16–19^ as well as humans ^20,21^.

Recent rodent work identifies sensory cortical disinhibition as a candidate causal mechanism for long-term fear memory ^22–24^. Fear conditioning transiently disrupts excitation-inhibition (E/I) balance in sensory cortical circuits, which releases cue-responsive neurons from inhibitory control—possibly via superficial layer cholinergic activation of parvalbumin (PV+) interneurons— thereby increasing their excitability and incorporation into a memory trace; conversely, experimentally blocking disinhibition prevents fear memory formation ^22–24^.

In humans, of particular relevance to a sensory cortical disinhibition account, fear conditioning consistently potentiates alpha-frequency event-related desynchronization (αERD; a stimulus-evoked reduction in α-band (8-12 Hz) power), selectively for conditioned threat versus safety stimuli (CS+ vs. CS-) ^25–27^. Alpha oscillations modulate sensory cortical inhibition ^28–33^, and occipital αERD is thought to reflect a release from inhibition, indexing visual cortical disinhibition ^31,32^. Accordingly, conditioning-potentiated αERD offers a plausible human neural signature of sensory cortical disinhibition during fear memory formation.

Disinhibition has emerged as a fundamental motif in neural processing, with maladaptive disinhibition and related E/I imbalance increasingly recognized as core pathophysiological features across a broad spectrum of psychiatric disorders ^34–36^. Thus, identifying sensory cortical disinhibition as a mechanism for long-term fear memory carries profound translational implications: exaggerated disinhibition could precipitate and perpetuate anxiety-related disorders whereas modulation of this process may serve as a therapeutic intervention.

Here, we directly tested the mechanistic role of sensory cortical disinhibition in human fear memory by experimentally counteracting it via high-definition (HD) α-frequency transcranial alternating-current stimulation (HD α-tACS) over early visual cortex during fear conditioning. Because α-tACS reliably augments α oscillations ^37–41^ and suppresses visual cortical activity ^42^, it affords a principled means to prevent CS+-potentiated occipital αERD and associated disinhibition during memory formation. We tracked fear memory development across its key stages (early formation, consolidation, and long-term retention) using transformer decoding of EEG and fMRI responses at 15 min, 24 h, and 7 days post-conditioning (**Fig. 1**). We hypothesized that α-tACS would blunt CS+-potentiated αERD and selectively disrupt fear memory consolidation and retention, thereby establishing sensory cortical disinhibition as a necessary mechanism for long-term fear memory in humans.

**Figure 1.**
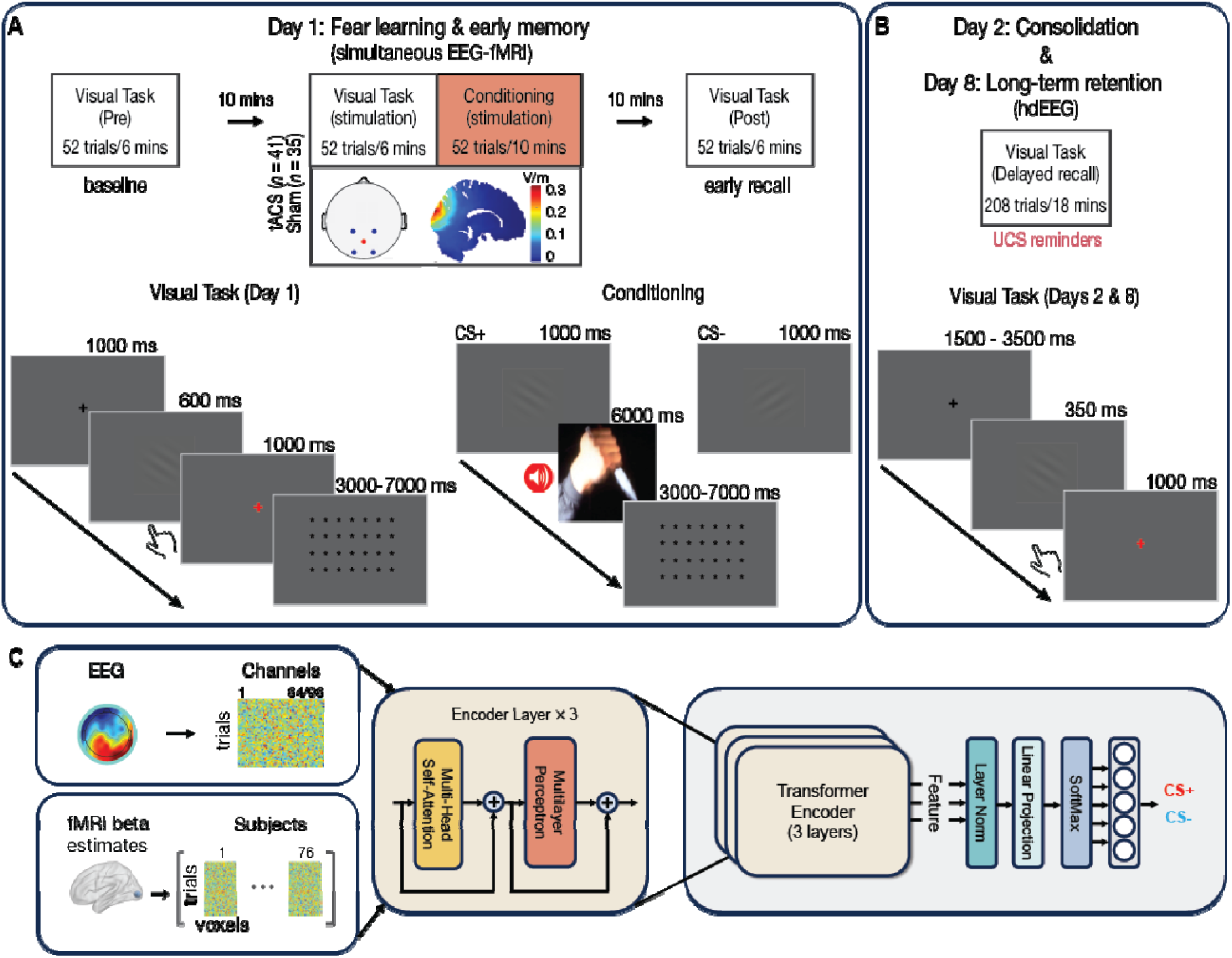
Experimental Paradigm and Analytical Design. (A) Day 1: Fear learning and early formation. Participants underwent fear conditioning during HD α-tACS (*n* = 41) or sham (*n* = 35). A visual task (VT) was administered before (VT-Pre), during stimulation (VT-Stim), and 15 min after conditioning (VT-Post) with simultaneous EEG-fMRI. In conditioning, one Gabor orientation (CS+) was paired with aversive audiovisual unconditioned stimuli (UCS; 50% reinforcement). In the VT, participants responded to a rare red crosshair following CS+ or CS- to maintain attention. HD α-tACS (4 × 1 montage, posterior midline) was delivered during VT-Stim and conditioning; finite-element modeling confirmed maximal field intensity in occipital visual cortex (0.34 V/m; cuneus, MNI 16, −93, 24). **(B) Days 2 & 8: Consolidation & Long-term retention.** A subsample (Active *n* = 23; Sham *n* = 18) returned for the VT (with 16 intermixed, unpaired UCS trials to limit extinction) during high-density EEG (hdEEG) at 24 h and 7 d post-conditioning. **(C) Transformer-based neural decoding.** Separate transformer models multi-head self-attention were trained on EEG and fMRI data to classify CS+ vs. CS-. Fear memory traces were indexed by the increase in decoding accuracy from pre- to post-conditioning indexed.

## Results

### α-tACS inhibits visual cortical activity

On Day 1, participants first performed a simple visual task (VT-Pre), where a CS was followed by a white (67 probability) or red (23% probability) crosshair, with a button press required for the red crosshair. Participants were then randomized to receive α-tACS (*n* = 42) or sham stimulation (*n* = 39) while repeating the visual task (VT-Stim). An fMRI double contrast (VT-Stim − VT-Pre, Active − Sham) validated the efficacy of our α-tACS protocol for inhibiting visual cortical activity (peak: x, y, z = 18, −68, 10; Z = 3.98, FDR *p =* 0.040 SVC), wherein the Active (but not the Sham) group showed reduced V1/V2 activation in VT-Stim (vs. VT-Pre; **Fig. 2A**). EEG data during VT-Stim and conditioning (below) were excluded from analysis due to stimulation artifacts.

**Figure 2.**
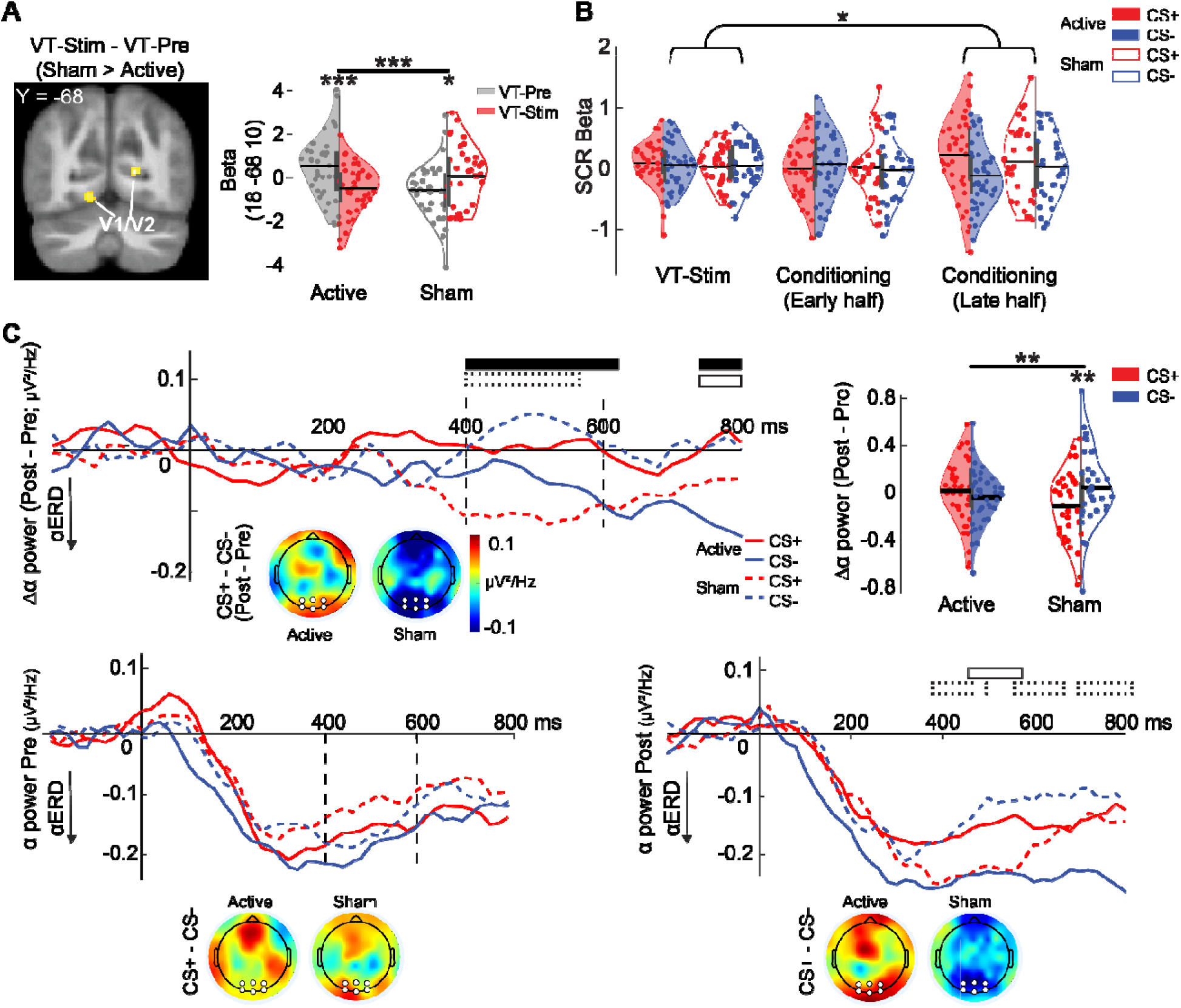
Experimental Validation. **(A) HD α-tACS induced visual cortical inhibition.** fMRI responses during VT-Stim (vs. VT-Pre) were reduced in the Active (vs. Sham) group. *Left*: Statistical parametric map (SPM; *p <* 0.005 uncorrected). *Right*: Peak voxel beta estimates from V1 (*Z* = 3.98, FDR *p =* 0.040 SVC). **(B) Fear conditioning heightened SCR.** SCR to CS+ versus CS-increased during the second half of fear conditioning, regardless of group. **(C)** α**-tACS blocked conditioning-potentiated** α **desynchronization.** Occipital α power (and difference) waveforms are shown (negative values indicate αERD). *Top*: Differential (VT-Post − VT-Pre) α power revealed robust CS+-potentiated αERD in the Sham but not Active group. Black horizontal bars denote contiguous time bins with significant Group-by-CS effects (FDR-corrected); bars with solid or dotted borders mark time bins with significant CS effects within Active and Sham groups, respectively (FDR-corrected). Topographical maps depict spatial distributions of differential αERD (400-600 ms), highlighting occipital channels (white dots = channels included in analysis). Violin plots display significant Group-by-CS interaction (400-600 ms): Sham exhibited potentiated αERD for CS+ (vs. CS-) at Post (vs. Pre); Active showed no effect; black horizontal line/grey box = mean/SEM. *Bottom:* Baseline (VT-Pre) αERD was equivalent across CS and groups (left). Post-conditioning (VT-Post), Sham exhibited the expected CS+-potentiated αERD, whereas Active showed a blunted/reversed pattern (right). \**p <* 0.05, \*\**p <* 0.01, \*\*\**p <* 0.001.

### Conditioning induces fear responses

Following the VT-Stim, participants underwent 10-minute fear conditioning alongside uninterrupted stimulation. A three-way rANOVA (Phase: VT-Stim/Early half/Late half of conditioning; CS, Group) on skin conductance responses (SCR β values) showed a Phase-by-CS interaction (*F =* 3.05, *p =* 0.051) but no Group main or interaction effects (*p*’s > 0.219; **Fig. 2B**). Follow-up tests revealed augmented SCR to CS+ versus CS-during late, but not early, half of conditioning (vs. VT-Stim; late: *t =* 2.54, *p =* 0.013; early: *t =* 0.31, *p =* 0.76). These results confirmed successful fear conditioning and indicated that α-tACS did not impair acquisition.

### α-tACS blocks CS+-potentiated α desynchronization

After a 10-minute rest inside the scanner (resulting in a ∼15-min post-conditioning window for early memory to form), participants completed a post-conditioning visual task (VT-Post). We found that α-tACS counteracted conditioning-induced αERD potentiation. A three-way rANOVA (Phase: Pre/Post, CS, Group) on EEG αERD (i.e., α power reduction following CS onset) revealed a significant three-way interaction (*F =* 8.56, *p =* 0.004; **Fig. 2C**, top row). Follow-up two-way (Phase, CS) rANOVAs showed a Phase-by-CS interaction in the Sham (*F =* 7.81, *p =* 0.008), but not Active group (*F =* 1.49, *p =* 0.234). Specifically, at VT-Pre, both groups exhibited equivalent αERD to CS+ and CS-(**Fig. 2C**, bottom left) but diverged at VT-Post with greater αERD to CS+ than CS-in Sham participants (*t =* −2.91, *p =* 0.006), while an opposite effect appeared in Active participants (*t =* 2.37, *p =* 0.023; **Fig. 2C**, bottom right). This suggests that α-tACS overrode CS+-potentiated α desynchronization. Furthermore, bin-by-bin analyses (FDR-corrected) localized this effect to 400-620 ms and 740-840 ms post-stimulus, similar to windows identified for previously reported CS+-potentiated αERD ^25–27^. These results thus replicated CS+-potentiated αERD and, critically, validated that our targeted α-tACS blocked this disinhibition.

### α-tACS of visual cortex disrupts fear memory traces across memory stages

With our conditioning and tACS manipulations validated, we sought to identify fear memory traces based on transformer decoding of EEG-fMRI responses to the CS. Decoding accuracy increases from pre-conditioning to 15 min, 24 h, and 7 days post-conditioning indexed fear memory traces at early formation, consolidation, and long-term retention, respectively. We then compared decoding accuracy increases between the Active and Sham groups to test the hypothesis that α-tACS of visual cortex would disrupt these fear memory traces.

### Fear Memory Early Formation (15 min)

#### Temporal profiles of CS decoding (EEG traces)

Using 64-channel EEG, for each participant, we derived bin-by-bin (20-ms resolution) CS decoding accuracy across 200-ms pre-stimulus and 600-ms stimulus presentation periods at VT-Pre and VT-Post (**Fig. 3A**). Supporting the sensitivity and specificity of our transformer decoding, baseline CS decoding hovered around chance (0.5) during the pre-stimulus interval, rising significantly (though modestly) at 40-60 ms post-stimulus and throughout the entire epoch (FDR *p <* 0.05; collapsed across groups). For hypothesis-testing, an rANOVA (Phase: Pre/Post, Group) on decoding accuracy indicated a Phase-by-Group interaction emerging at 60-80 ms (*F =* 22.95, *p <* 0.001) and throughout the epoch (FDR *p <* .05). Specifically, post-conditioning increases in decoding accuracy were markedly weaker in the Active than Sham group, albeit significant in both groups (Active: *t =* 2.36.98, *p =*0.023; Sham: *t =* 7.95, *p <* .001). Therefore, reliable fear traces emerged in EEG at early memory formation, supporting fear reactivation at an initial sensory stage (40-60 ms), but α-tACS of visual cortex attenuated this plasticity.

**Figure 3.**
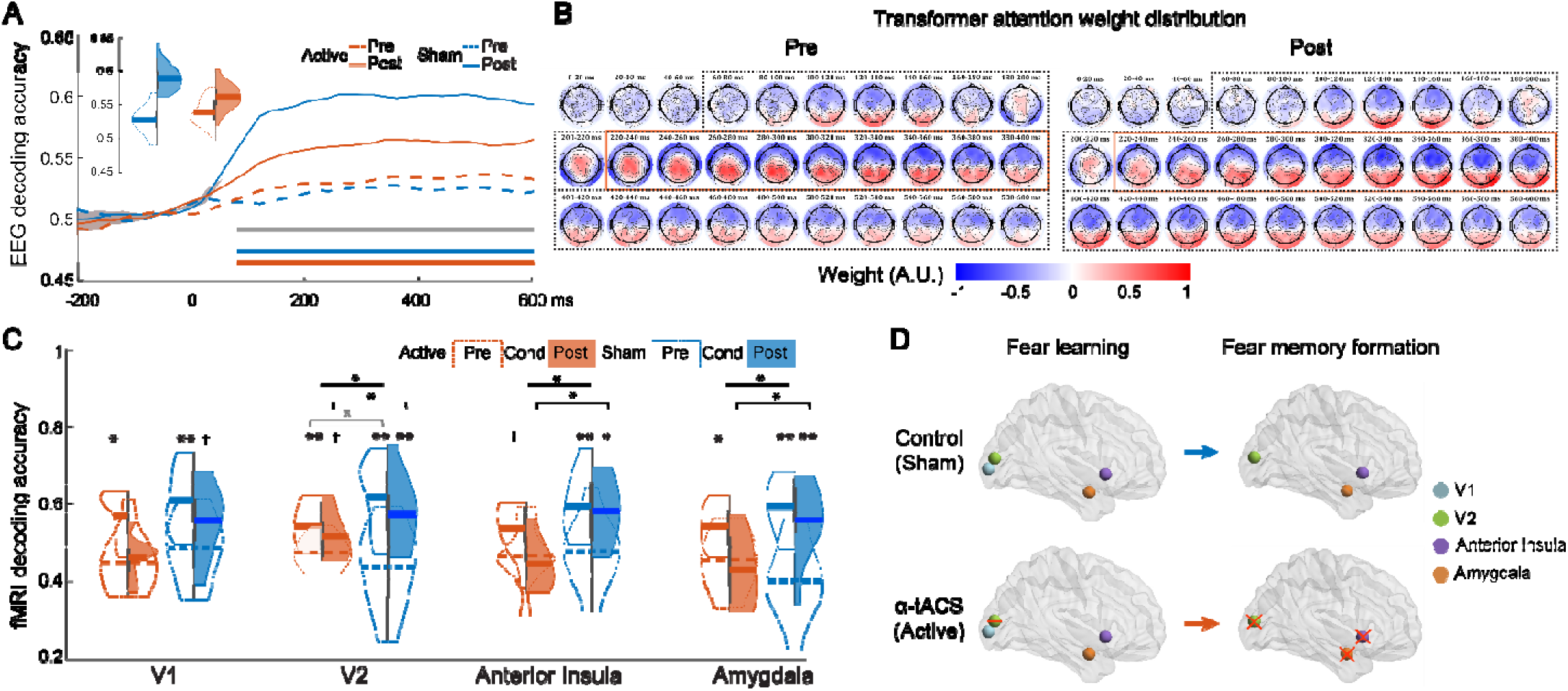
HD α-tACS of visual cortex disrupted EEG and fMRI traces of early memory formation (Day 1). **(A): EEG decoding accuracy.** CS decoding improved from VT-Pre to VT-Post (15 min) in both Active (orange) and Sham (blue) groups (horizontal bars: contiguous significant time bins, FDR-corrected). This effect was stronger in Sham (gray bar: significant Phase-by-Group effects, FDR-corrected). Shaded ribbon = SEM. Inset: violin plots of decoding accuracies at VT-Pre (light) and VT-Post (dark). **(B) Transformer attention weight maps.** Spatiotemporal dynamics of attention weights (channel contributions) indicated occipital dominance during early CS decoding, followed by a broadened central distribution, before returning to occipital channels. Black dotted boxes: time bins with above-chance decoding; red boxes: increased occipital contributions post-conditioning. Maps were comparable across groups and are shown averaged (see **Extended Data Fig. 1** for group-specific maps). **(C) fMRI decoding accuracy.** All visual and limbic ROIs showed decoding improvement during conditioning (vs. VT-Pre) in both groups. At VT-post, Sham alone showed decoding improvement in V2, anterior insula, and amygdala, but not V1. **(D) Graphical summary.** V1 showed decoding improvement during conditioning alone. V2, anterior insula, and amygdala exhibited decoding improvement in both conditioning and early memory formation, with the latter selectively preserved in Sham. Red bar: attenuation by α-tACS; red cross: elimination by α-tACS. * = *p <* 0.05; ** = *p <* 0.01, † = *p <* 0.1.

Scalp topographical maps of transformer attention weights (i.e., channel contributions) further revealed time-resolved spatial dynamics of fear reactivation (**Fig. 3B**). Bin-by-bin topographical analyses of variance (TANOVA; Phase, Group) isolated a Phase effect throughout 220-400 ms (FDR *p* = 0.003), reflecting a weight shift towards occipital channels from pre- to post-conditioning, with no effect of Group or Group-Phase interaction (**Extended Data Fig. 1**). These topographical changes underscored visual cortical involvement in fear memory formation.

#### CS decoding in the sensory-limbic circuit (fMRI traces)

Using simultaneously acquired fMRI, we next examined CS decoding in sensory-limbic fear regions, focusing on early visual areas (V1/V2) and limbic areas strongly implicated in human fear conditioning: anterior insula and amygdala ^3,43^ (**Fig. 3B&C**). An rANOVA (Phase: VT-Pre/Conditioning/VT-Post, Group) on V1 decoding showed an effect of Phase (*F =* 14.13, *p <* 0.001), driven by decoding accuracy increase from VT-Pre to conditioning (*t =* 4.81, *p <* 0.001) and a trend from VT-Pre to VT-Post (*t =* 1.73, *p =* 0.10), with no Phase-by-Group interaction (*p =* 0.386). Thus, V1 exhibited reliable fear learning traces but only weak fear memory traces. Notably, α-tACS did not affect fear learning traces in V1.

By contrast, a similar rANOVA on V2 decoding showed a Phase-by-Group interaction (*F =* 4.48, *p =* 0.025). From VT-Pre to conditioning, both groups exhibited decoding accuracy increase (Sham/Active *t =* 4.16/3.17, *p =* 0.002/.011), an effect that was stronger in the Sham (vs. Active) group (*t =* 2.33, *p =* 0.037). From VT-Pre to VT-Post, this decoding improvement persisted in the Sham group (*t =* 4.01, *p =* 0.003) but reduced to a trend in the Active group (*t =* 2.20, *p =* 0.055), resulting in a significant group difference (*t =* 2.52, *p =* 0.025). Thus, V2 exhibited reliable traces for both fear learning and early memory, which nonetheless were vulnerable to α-tACS, particularly during early memory formation.

The limbic regions exhibited patterns paralleling V2. In anterior insula, a Phase-by-Group interaction (*F =* 4.31, *p =* 0.026) indicated intact learning traces but impaired memory traces under α-tACS: decoding accuracy increased from VT-Pre to conditioning across participants (*t =* 4.17, *p =* 0.001), with no group difference (*p =* .282), but only the Sham group maintained this gain at VT-Post (*t =* 2.78, *p =* 0.022; Active: *p =* 0.473; group difference: *t =* −2.54, *p =* 0.021). The amygdala mirrored this dissociation (Phase × Group: *F =* 4.75, *p =* 0.016): during conditioning, both groups showed decoding accuracy increase (*t =* 4.75, *p <* 0.001; with a trend towards a group difference: *t =* −1.84, *p =* 0.082), while at VT-Post, only the Sham group retained this improvement (*t =* 2.87, *p =* 0.018; Active: *p =* 0.448; group difference: *t =* −2.86, *p =* 0.012).

Overall, Day 1 fMRI results revealed a convergent profile across secondary visual cortex and limbic regions: decoding improvement emerged during conditioning and persisted in early memory formation, but the latter was selectively disrupted by α-tACS (**Fig. 3D**). V1, by contrast, expressed improvement during conditioning alone, which was insensitive to stimulation. These findings suggest that fear learning engages a broad sensory-limbic network, whereas early fear memory formation recruits specifically secondary sensory cortex and limbic structures. Critically, consistent disruptions of EEG-fMRI fear memory traces under targeted α-tACS of visual cortex isolate sensory cortical disinhibition as a required mechanism for human fear memory formation.

### Fear Memory Consolidation and Long-term Retention (24 hr & 7 days)

A subsample of 41 participants (Active *n* = 23; Sham *n* = 18) returned at 24 hr and 7 days for another VT (VT-Day 2 and VT-Day 8) while 96-channel hdEEG was acquired. The decoding procedure was virtually identical to Day 1 (for each participant, 20-ms resolution, across the 200-ms pre-stimulus and 360-ms stimulus presentation periods).

#### Consolidation (24 hr)

Bin-by-bin rANOVAs (Phase: Pre/Day 2, Group) on decoding accuracy revealed a Phase-by-Group interaction (*F =* 4.75, *p =* 0.036), emerging at 40-60 ms and persisting to 200 ms (FDR *p <* 0.05; **Fig. 4A**, left). Follow-up *t*-tests (VT-Day 2 vs. VT-Pre) showed decoding improvement in the Sham group (FDR *p <* 0.05), but no change in the Active group at any time bin (*p*’s > .099). Topographical maps of transformer attention weights revealed a significant Group effect across the entire epoch (FDR *p* < 0.001; **Fig. 4A**, center): the Sham group preserved the spatiotemporal pattern at Day 1 Post, which was however lost in the Active group, paralleling the lack of decoding change in this group (**Extended Data Fig. 1**). These results revealed EEG traces for fear memory consolidation, engaging early sensory processing in fear reactivation, which were nonetheless abolished by targeted α-tACS.

**Figure 4.**
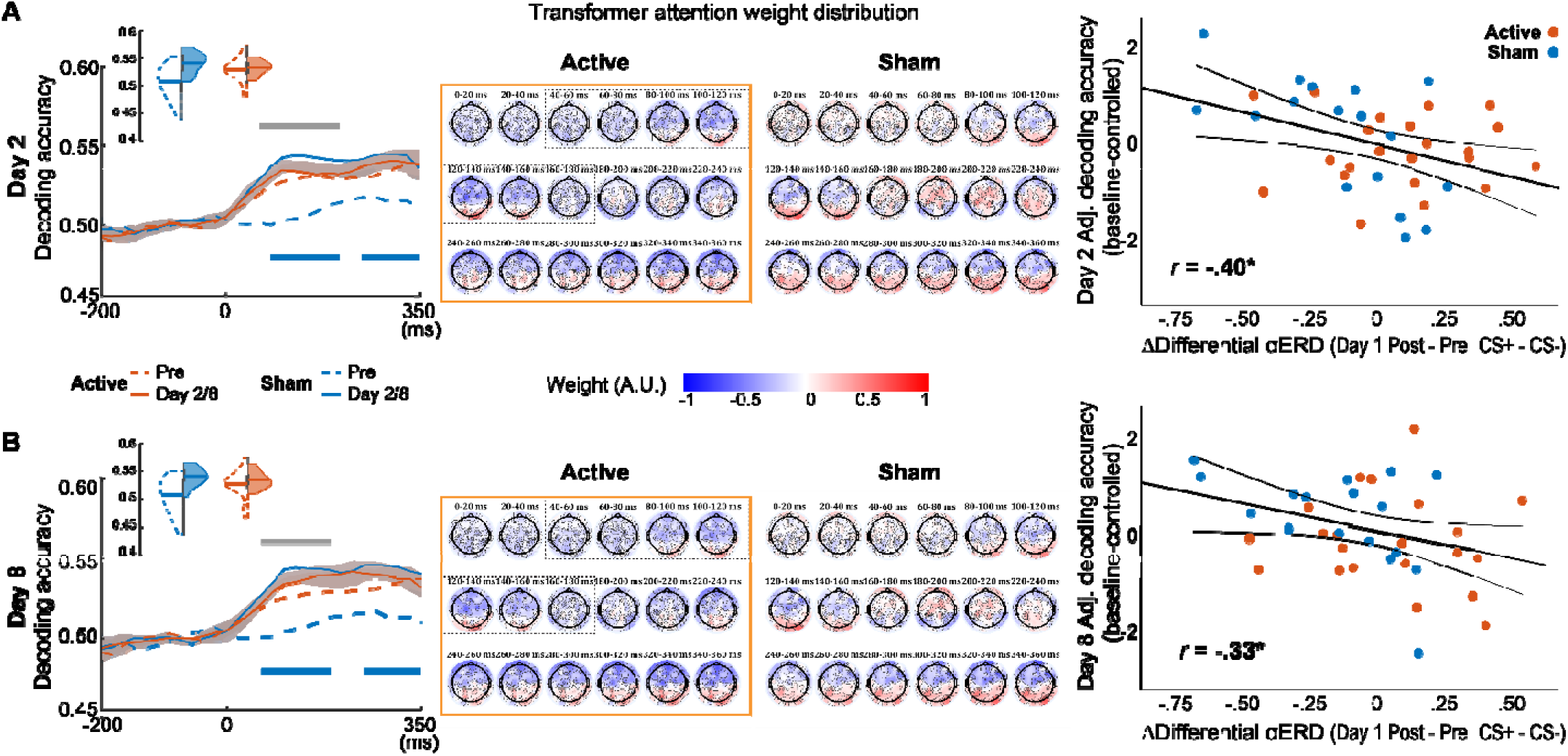
HD α-tACS of visual cortex disrupted fear memory consolidation and long-term retention. (**A**) **Consolidation** (**Day 2**) and (**B**) **Long-term retention** (**Day 8**). *Left*: CS decoding improved from VT-Pre on both days in the Sham (blue), but not Active (orange), group. Gray horizontal bars indicate contiguous time bins with significant Group effects (tACS disruption of decoding improvement; FDR-corrected; Day 2: 40-200 ms; Day 8: 40-180 ms). Blue horizontal bars denote significant decoding improvement within Sham (FDR-corrected; Day 2: 60-200 ms; Day 8: 40-180 ms). Shaded ribbon = SEM. Insets: violin plots of decoding accuracies at VT-Pre (light) and VT-Day2/Day8 (dark) for both groups. *Center*: Transformer attention weight maps. Dotted boxes indicate time bins with significant Phase-by-Group effects; red boxes denote Group effects spanning the entire epoch; Sham preserved baseline spatiotemporal dynamics— early occipital dominance, transient broad distribution, and later return to occipital dominance— which were absent in Active. *Right*: Disinhibition-memory association. Baseline-adjusted (adj.) decoding accuracy (Day 2 and Day 8, controlling for Day 1 decoding) was negatively predicted by CS+-potentiated αERD. \**p <* 0.05.

#### Long-term retention (7 days)

Similar bin-by-bin rANOVAs (Phase: Pre/Day 8, Group) on Day 8 again revealed a Phase-by-Group interaction (*F =* 6.828, *p =* 0.013), emerging at 40-60 ms and persisting to 180 ms (**Fig. 4B**, left). Follow-up *t*-tests (Day 8 vs. Pre) showed significant decoding improvement in the Sham group (FDR *p <* 0.05), but not in the Active group (*p*’s >.224). Transformer attention maps again revealed a Group effect throughout the epoch (FDR *p* < 0.001), with the Sham group recapitulating Day 2 spatiotemporal dynamics that were again lost in the Active group (**Fig. 4B**, center; **Extended Data Fig. 1**). These results paralleled Day 2 findings, revealing long-term fear memory traces grounded in early sensory processing but eliminated by targeted α-tACS.

### Linking CS+-potentiated α desynchronization to long-term fear memory

To establish a direct link between conditioning-induced visual cortical disinhibition and fear memory, we tested whether CS+-potentiated occipital αERD predicted the strength of memory traces at the three key stages. In three separate multiple regressions (baseline/VT-pre decoding entered as covariate), differential α power (VT-Post − VT-Pre; CS+ − CS-) negatively predicted decoding accuracy at 24 hr (*r* = −.39, *p =* .015; **Fig. 4A**, right) and 7 days (*r* = −.33, *p =* .042; **Fig. 4B**, right), but not at 15 min (*r* = −.16, *p =* .166; also see **Extended Data Table 1**). These results underscore a close association between sensory cortical disinhibition and fear memory consolidation and long-term retention.

## Discussion

By integrating multimodal neuroimaging, transformer-based decoding, and brain stimulation, we uncover cohesive spatiotemporal traces of fear memory embedded within sensory-limbic circuits across early formation (15 min), consolidation (24 h), and long-term retention (7 days). Fear conditioning rapidly enhanced occipital α desynchronization, indexing CS+-related visual cortical disinhibition; importantly, this disinhibitory effect predicted the strength of subsequent memory consolidation and retention. We further establish a causal role for this visual cortical process using HD α-tACS of the visual cortex. This targeted stimulation suppressed V1/V2 activation and abolished CS+-potentiated α desynchronization, confirming the engagement of blocking visual cortical disinhibition. Critically, this principled blockage of visual cortical disinhibition disrupted fear memory development, particularly, consolidation and long-term retention, while leaving initial fear acquisition intact.

Together, these findings demonstrate that human fear memory engages distributed plasticity across sensory and limbic regions. Importantly, they identify sensory cortical disinhibition as a causal, memory-stage-specific mechanism, underlying memory consolidation and retention (but not acquisition). More broadly, our results highlight sensory cortex as a key locus of enduring fear memory representations (or “schemas”) in humans ^14,44^ and position it as a mechanistically-grounded target for modulating persistent fear in anxiety- and trauma-related disorders, complementing existing limbic-prefrontal-focused models ^45^.

Compared to non-human studies, human evidence regarding the neural basis of long-term fear memory is limited. Here, fMRI-EEG testing at three key memory stages provides novel spatiotemporal insights into the neural dynamics underpinning long-term fear memory in humans. Key limbic regions (i.e., amygdala and anterior insula) showed an expected plasticity during fear conditioning and early recall, affirming their roles in fear acquisition and early memory consolidation ^2,3^. Notably, while amygdala plasticity often elude detection in univariate fMRI analysis ^3^, our use of AI-based multivariate analysis captured it robustly. In visual cortex, decoding improvement arose in both V1 and V2 during fear conditioning but persisted only in V2 at recall, a finding that aligns with prior animal ^13,17–19^ and human ^7,20^ evidence highlighting secondary sensory cortex in fear memory.

Our EEG-EEG decoding tracks fear memory traces across formation, consolidation, and long-term retention and provides pivotal temporal insights. Scalp topography of decoding weights reveals time-resolved spatial dynamics of fear memory reactivation, consistent across all three retests, characterized by an early occipital dominance (starting at 60-80 ms), coinciding with V1/V2 fear response latencies in primates ^46,47^, which then gives rise to a broader, central distribution (180-300 ms). This spatiotemporal dynamic strengthens prior accounts of the time course of fear perception ^14,48^, aligning with a recent finding of cross-species temporal progression in response to emotionally salient sensory cues ^49^. Notably, conditioning engendered a decoding-weight shift towards occipital channels that persisted at 24 hr and 7 days, underscoring visual cortical contributions to long-term fear memory.

Targeted HD α-tACS of visual cortex impacted all fMRI/EEG memory traces, including the aforementioned spatiotemporal dynamics. Consistent with prior reports ^42^, our α-tACS procedure suppressed visual cortical responses in VT-stim, producing visual cortical inhibition. Critically, this further blocked CS+-potentiated αERD (an index of sensory disinhibition/excitation ^31–33^), which was nonetheless evident in our Sham group as previously observed during both late fear conditioning and early fear recall ^25–27^. Importantly, the magnitude of CS+-potentiated αERD predicted the strength of long-term memory traces (at 24 hr and 7 days, albeit not shortly after), emphasizing a link between visual cortical disinhibition and processes pivotal for enduring fear memories—consolidation and retention. Notably, α-tACS did not affect CS decoding improvement or CS+-related SCR increase during conditioning. These observations suggest that sensory cortical disinhibition is not critical for the aspect of fear learning and rule out the alternative explanation that the disrupted memory traces stemmed from impaired fear learning.

fMRI decoding confirmed sensory-limbic engagement during fear learning and early recall, with its disruption by α-tACS during early recall indicating that this involvement depends on visual cortical disinhibition. Although EEG memory traces were markedly attenuated by α-tACS, they remained detectable at early recall, likely due to EEG’s whole-scalp coverage capturing the global fear network (beyond sensory-limbic circuits) as well as its superior reliability and sensitivity (relative to fMRI) ^50^. Notably, despite these technical advantages, by 24 hr and 7 days, EEG memory traces were no longer evident in the Active group, underscoring stage-specific (consolidation and retention) role of sensory cortical disinhibition in fear memory.

Together, these multimodal fMRI-EEG findings coalesce to support models emphasizing sensory cortex as a key (extra-limbic) substrate for fear memory, and critically, establish sensory cortical disinhibition as a causal mechanism for enduring fear memories in humans. Clinically, these results add to the growing recognition of E/I imbalance as a core feature of anxiety-related disorders. Furthermore, the ability of α-tACS to target and block this mechanism opens a potential new avenue for therapeutic intervention, particularly in thwarting pathological fear central to anxiety- and trauma-related disorders.

## Methods

Eighty-one healthy adults with normal or corrected-to-normal vision were recruited in the study. Participants reported no history of head injury, psychiatric or neurological disorders, or current use of psychotropic medication. Absence of current psychiatric disorders was determined through the Structured Clinical Interview for DSM-V. All participants abstained from alcohol (24 h) and caffeine (4 h) prior to testing. Following informed consent (Florida State University IRB approved), participants were randomly assigned to sham (*n* = 39) or active stimulation (*n* = 42). Five were excluded due to motion, poor EEG quality, or equipment malfunction, yielding a final sample of 76 (Active *n* = 41; Sham *n* = 35; 53 female; mean age = 23.0 ± 6.2 years, range 18-46). A subsample of 41 (Active *n* = 23; Sham *n* = 18; 25 female; mean age = 21.5 ± 4.8 years, range = 18-38) further underwent recall testing on Day 2/Day 8. Two (Active *n* = 1; Sham *n* = 1) participated only on Day 2, and two (Active *n* = 1; Sham *n* = 1) only on Day 8, resulting in a final retest sample of 39 (Active *n* = 22; Sham *n* = 17) on either day.

### Experimental Paradigm

The experiment employed a mixed factorial design with Conditioned Stimulus (CS+/CS-), Phase (Pre, Conditioning, Post, Day 2, Day 8), and Group (Active/Sham) as factors. As shown in **Fig. 1**, a visual task (VT) was administered at baseline (VT-Pre), during α-tACS/sham stimulation (VT-Stim), and at three post-conditioning time points (15 minutes, 24 hours, and 7 days later; VT-Post, VT-Day 2, VT-Day 8). The conditioning phase immediately followed VT-Stim and was conducted while α-tACS/sham stimulation continued. A 10-minute resting interval was then included, separating VT-Post from the conditioning phase by approximately 15 minutes to permit early fear memory formation. On Day 1, simultaneous EEG-fMRI was recorded throughout the experiment, while high-density EEG (hdEEG) was collected on Days 2 and 8.

#### Stimuli

Unconditioned stimuli (UCS) were adapted from our previous work ^21^ and consisted of bimodal audiovisual presentations combining images from the International Affective Picture System (IAPS) ^51^ with sounds from the fear subset of human affective vocalizations ^52^. CS were centrally presented Gabor patches (sinusoidal gratings with Gaussian envelope; 5.5° × 5.5° visual angle), tilted either 30° left or right of vertical to serve as CS+ and CS-(counterbalanced across participants). The Gabor patches were achromatic, with spatial frequency of 2.2 cycles/°, luminance contrast of 6.9% Michelson contrast, and luminance range 20.16-23.14 cd/m², presented against a gray background (21.65 cd/m²). These CS were used in both conditioning and the VT task. They were displayed using a high-precision MRI-compatible projector (PROPixx, VPixx Technologies) on Day 1 and on a CRT monitor on Days 2 and 8.

#### Fear Conditioning

During conditioning, Gabor patches were presented for 1 s in pseudorandom order (no more than three consecutive CS of the same type). A total of 50 CS+ trials were delivered, with 25 (50% reinforcement) followed by a 6-s UCS presentation, along with 25 CS-trials. Inter-trial intervals consisted of a black-star screen presented for 3000-7000 ms. To ensure attention to the stimuli, participants were instructed to respond to a red crosshair via button pressing on 6 unreinforced CS+ or CS-trials (12% probability).

#### Visual Task (VT)

During VT-Pre, VT-Stim, and VT-Post on Day 1, each trial began with a central black crosshair (1000 ms), followed by a CS+ or CS-(600 ms), and then a red or white crosshair (1000 ms). Participants responded via button press to the red crosshair (23% probability, evenly distributed between CS+ and CS-; **Fig. 1A**). Each stimulus was followed by a black-star screen (3000-7000 ms). Trials were presented in pseudorandom order (no more than three consecutive CS of the same type), with 52 trials per task (26 CS+, 26 CS-). To reduce extinction, VT-Post began with a UCS trial ^53^.

On Days 2 and 8, when only hdEEG was recorded, VT-Day 2/VT-Day 8 trial length was shortened: a black crosshair was presented for 1500-3500 ms, followed by a CS (350 ms) and then a red or white crosshair (1000 ms; **Fig. 1B**). A total of 208 trials were presented (104 CS+, 104 CS-), with 16 UCS trials randomly intermixed across the task to reduce extinction.

#### HD α-tACS

High-definition alpha-frequency transcranial alternating current stimulation (HD α-tACS) was administered using an MR-compatible system (Soterix Medical, New York, NY), following procedures from our prior studies ^40,41^. Participants were randomly assigned to receive either active stimulation (2 mA sinusoidal, 10 Hz) targeting the occipital visual cortex or sham stimulation as a control. The stimulation montage consisted of a central anode surrounded by four cathodes in a 4 × 1 high-definition electrode configuration (**Fig. 1A**). Stimulation was delivered for 20 minutes, coinciding with the visual task (VT-Stim) and fear conditioning.

### Data Acquisition and Preprocessing

#### EEG

On Day 1, as in our prior study ^40^, simultaneous EEG-fMRI data were acquired using an MR-compatible 64-channel EEG system (Brain Products GmbH, Germany), sampled at 5000 Hz, synchronized with the scanner clock, and filtered online at 0.05-200 Hz. Concurrent electrocardiogram (ECG) was recorded with a shielded electrode for cardioballistic artifact removal. MR- and cardioballistic-related artifacts were corrected offline using BrainVision Analyzer 2.0 (Brain Products GmbH). On Days 2 and 8, hdEEG was collected with a 96-channel actiCAP system (Brain Products GmbH, Germany) including electrooculogram (EOG) channels. All EEG data were referenced to the average of all electrodes. Artifact detection and correction were performed using the Fully Automated Statistical Thresholding for EEG artifact Rejection (FASTER) algorithm implemented in EEGLAB ^54^. Data were downsampled to 250 Hz, bandpass filtered (0.1-40 Hz), and epoched from −400 to 1000 ms relative to CS onset. Epochs exceeding ±3 SDs in amplitude range, variance, or deviation were rejected. Independent component analysis (ICA; Infomax algorithm; ^55^) was applied to remove artifactual components (e.g., muscle activity, blinks, saccades, electrode pops). Remaining deviant channels within cleaned epochs were interpolated using spherical spline interpolation in EEGLAB.

#### EEG Event-Related Desynchronization (ERD)

ERD on Day 1 was estimated by first epoching the EEG data from −400 to 1000 ms relative to CS onset, followed by subtraction of the mean event-related potential (ERP). α-band power (8-12 Hz) was then extracted from six occipital electrodes (Oz, O1, O2, POz, PO3, PO4) using the multitaper spectral estimation technique within 200-ms windows stepped in 20-ms increments. α power was normalized relative to the pre-stimulus window of each trial ^27^ and subsequently averaged across channels and trials for each CS type. This yielded post-stimulus αERD waveforms, and for hypothesis testing, we focused on αERD during the 400-600 ms post-stimulus interval, consistent with prior studies ^25–27^.

#### EEG Decoding Analysis

Decoding analyses followed the procedures in our prior work ^56^. To reduce noise and enhance robustness, five consecutive EEG data bins (non-overlapping) were averaged, yielding a smoothed timeseries with a temporal resolution of 20 ms. Decoding was performed on data from −200 to 600 ms (Day 1) and −200 to 350 ms (Days 2 and 8) relative to stimulus onset. The pre-stimulus period served as baseline, and the stimulus presentation durations defined the windows of interest, i.e., 0-600 ms on Day 1 and 0-350 ms on Days 2/8.

As illustrated in **Fig. 1C**, at each time bin, CS+ and CS-trials were treated as independent observations, with the EEG channels (64 on Day 1; 96 on Days 2 and 8) serving as the feature set. This yielded a data matrix per participant per VT session, which was then entered into a three-layer transformer encoder based on a visual transformer (ViT) architecture. The encoder consisted of three attention blocks, with output from the final (third) block passed through layer normalization, a linear projection, and lastly a SoftMax classifier.

Model performance was evaluated using five-fold cross-validation. For Day 1, training folds included 20 CS+/CS-trial pairs, with 6 pairs reserved for testing. For Days 2 and 8, training folds included 84, 84, 84, 84, and 80 pairs, with corresponding test folds of 20, 20, 20, 20, and 24 pairs, respectively. Each iteration ensured that all trials were tested exactly once. Decoding accuracy was computed per fold, then averaged across folds to yield participant-level accuracy at each time bin.

Finally, we derived EEG-channel-wise attention weights from the transformer at each time bin for each participant. These weights were normalized and averaged across participants to generate dynamic topographical attention maps, indicating the relative contribution of each channel to decoding decisions. We applied TANOVA ^57^ on the maps and extracted global map dissimilarity (GMD), defined as the root-mean-square of channel-wise differences between phases and groups. Simple and double contrasts were performed to test the effects of Phase, Group, and Phase-by-Group interaction based on nonparametric permutation. To control for multiple comparisons across time bins of the epoch, we applied temporal cluster-based permutation correction (cluster-corrected α = 0.05).

#### fMRI Acquisition and Preprocessing

On Day 1, as in our prior study ^40^, gradient-echo T2*-weighted images were acquired simultaneously with EEG on a 3T Siemens Prisma scanner using a 64-channel head coil with axial acquisition (TR/TE = 1800/22 ms; slice thickness/ga*p =* 1.8/0.45 mm; voxel size = 1.8 × 1.8 mm; GRAPPA acceleration factor = 2; multiband factor = 2). A high-resolution 3D-MPRAGE T1-weighted image (1 × 1 × 1 mm³) was also obtained. Initial preprocessing of anatomical and functional volumes was performed using fMRIprep ^58^, including spatial realignment, slice-timing correction, coregistration to T1, normalization to MNI space (MNI152Lin), and smoothing with a 6-mm FWHM Gaussian kernel.

#### fMRI Univariate Analysis

To validate tACS-induced visual cortical inhibition, preprocessed fMRI data from VT-Pre and VT-Stim were analyzed with a block-design general linear model (GLM) in SPM12. Each phase (onset of first trial to end of last trial) was modeled as a regressor convolved with a canonical hemodynamic response function (HRF) and its temporal and dispersion derivatives. Six motion parameters were included as regressors of no interest, with a high-pass filter (128 s) and AR(1) model applied. Subject-specific contrasts (VT-Stim − VT-Pre) were entered into between-group *t*-tests (Active vs. Sham). Small-volume correction (SVC, *p <* 0.005 FDR) was applied to right-lateralized V1 and V2, based on prior lab findings ^40,41^, and defined using SPM Anatomy Toolbox 2.2b ^59^.

#### fMRI Decoding Analysis

For hypothesis testing, preprocessed data from VT-Pre, VT-Stim, fear conditioning, and VT-Post Day 1 were modeled using a Least Squares All (LSA) GLM in SPM12. Each unreinforced CS+ or CS-trial was modeled as a separate regressor, while all reinforced CS-UCS trials were included as a single regressor, convolved with a canonical HRF. Motion parameters, high-pass filtering (128 s), and AR(1) correction were included. Beta coefficients for each CS trial were extracted from voxels within regions of interest (ROIs), concatenated across trials and participants to form group-level input matrices. These matrices were submitted to a three-layer transformer encoder for CS decoding. Encoder performance was evaluated with 10-fold cross-validation (Active: *n* = 41; Sham: *n* = 35), training on *n*\*42 trials and testing on *n*\*10 trials per fold, yielding decoding accuracies for statistical analysis.

#### Regions of Interest (ROIs)

For decoding of fear traces, we focused on regions important for fear processing, including bilateral V1, V2, amygdala, and anterior insula. V1 and V2 were defined using probabilistic maps at 50% threshold ^60^; the amygdala ROI was drawn from the Harvard-Oxford Atlas ^61^; and the anterior insula ROI from the Willard Functional Atlas ^62^.

#### Skin Conductance Response (SCR)

SCR was recorded simultaneously with EEG and fMRI using the Brain Products system. Data were downsampled to 250 Hz and processed using the SCR module of PsPM ^63^, which included bandpass filtering (0.1-5 Hz), further downsampling to 10 Hz, and participant-wise z-score transformation. The conservative bandpass filter was chosen to enhance sensitivity in modeling-based SCR analyses for simultaneous fMRI-SCR paradigms ^64^. A GLM based on the canonical SCR function was fit in PsPM for each participant, with regressors for CS+ (unreinforced), CS+ (reinforced), and CS-trials. To characterize the temporal dynamics of fear learning, the conditioning phase was divided into early and late halves as separate blocks in the GLM. Resulting β values for CS+ (unreinforced) and CS-trials were extracted for each participant and submitted to subsequent analyses.

### Statistical Analysis

We first validated (i) tACS-induced visual cortical inhibition, (ii) fear conditioning, and (iii) tACS suppression of conditioning-potentiated visual cortical disinhibition. For (i) visual cortical inhibition, beta estimates during VT-Stim versus VT-Pre were compared across groups, testing the contrast Sham (VT-Stim − VT-Pre) > Active (VT-Stim − VT-Pre). For (ii) fear conditioning, SCR β values were submitted to a repeated-measures ANOVA (rANOVA) with factors CS-type (CS+/CS-), Group (Active/Sham), and Phase (VT-Stim, early VT-Cond, late VT-Cond). For (iii) tACS suppression of cortical disinhibition, αERD was submitted to a three-way rANOVA with factors CS-type (CS+/CS-), Group (Active/Sham), and Phase (Pre vs. Post).

To test the key hypothesis of tACS disruption of neural plasticity related to fear memory, separate rANOVAs were conducted on EEG and fMRI decoding accuracy. For EEG, rANOVAs were performed at each time bin with factors Group (Active/Sham) and Phase (Pre vs. Post/Day 2/Day 8; *p <* .05, two-tailed), followed by multiple comparisons across time bins using cluster-based permutation in the permuco package (*n* = 5,000; threshold = 95th percentile). Significant interaction effects were followed up by within-group t-tests (*p <* 0.05, two-tailed). For fMRI, rANOVAs were conducted separately within each ROI, with factors Group (Active/Sham) and Time (VT-Pre, Conditioning, VT-Post). Significant interactions were followed up with within-group t-tests (VT-Pre vs. Conditioning; VT-Pre vs. VT-Post) to evaluate neural traces of fear learning and memory in each group (*p <* 0.05, two-tailed).

Finally, to examine whether tACS suppression of conditioning-induced cortical disinhibition predicted fear memory disruption, differential αERD (CS+ − CS-, Post Day 1 − Pre) was used as a predictor in multiple regression models, with Day 2 or Day 8 decoding accuracy as the outcome and baseline decoding accuracy included as a covariate.

## Competing Interest Statement

The authors declare no competing financial interests.

## Funding

This research is supported by the National Institutes of Health grants (R21MH126479 & R01MH132209 W.L.).

## Authors contributions

JAB and WL designed research; JAB, JJM, KC, and CC performed research; JAB, YM, YW, MD, CY, and WL analyzed data; JAB, YM, YW, CC, RJD, MRM, MD, CY, and WL wrote the paper.

## Data and materials availability

Source Data will be provided with this paper. Upon acceptance of the manuscript, data included in the study will be deposited in the lab’s GitHub repository (https://github.com/WenLiCssp).

## Code Availability

Upon acceptance of the manuscript, in-house code included in the study will be deposited in the lab’s GitHub repository (https://github.com/WenLiCssp).

All display items presented in the main manuscript can be reproduced from the data and code shared in the public repository.

---

**Extended Data Fig. 1.**
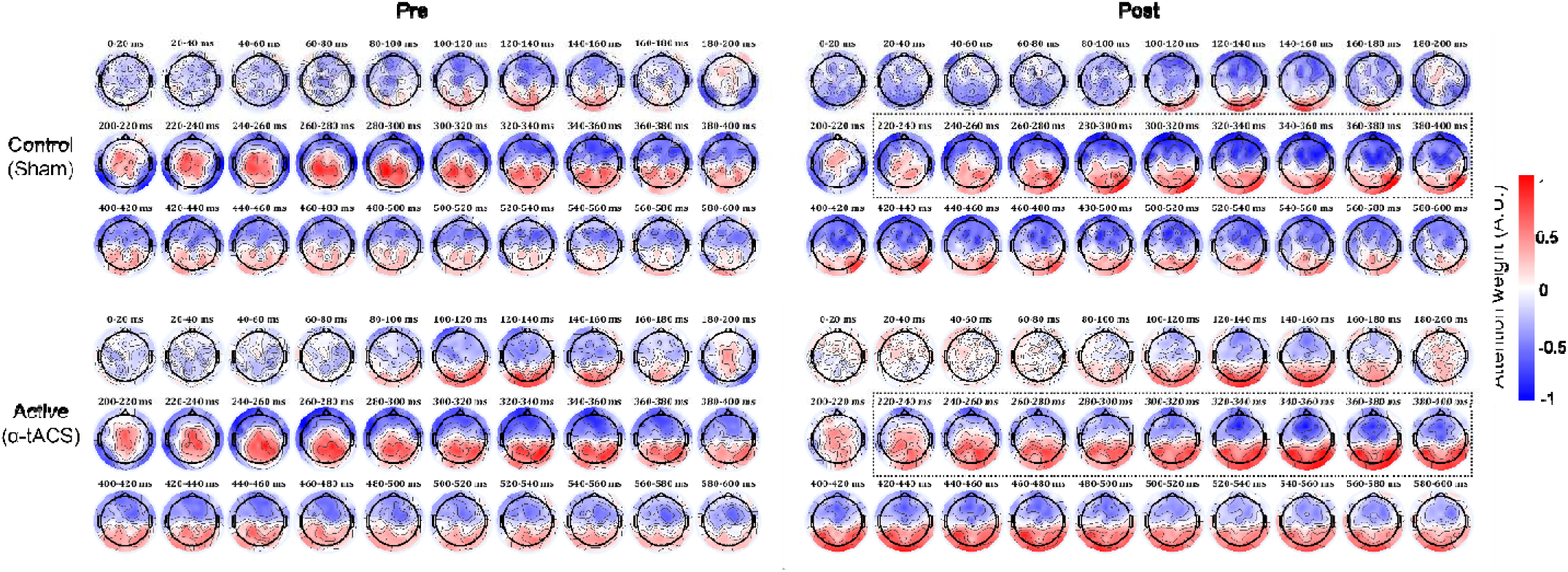
Transformer attention weight maps for Day 1. Topographical analysis of variance (TANOVA) ^57^ was used to quantify condition differences in attention-weight topographies using global map dissimilarity (GMD; root-mean-square of electrode-wise differences) computed for each time bin. Phase, Group, and Phase × Group effects were assessed with nonparametric permutation testing. No main effect of Group or Phase × Group interaction was detected (all *p*’s > 0.119). A significant main effect of Phase emerged at 220-400 ms (marked by the dotted boxes; FDR *p* = 0.003), reflecting a posterior shift in attention weights from Pre to Post.

**Extended Data Table. 1.**
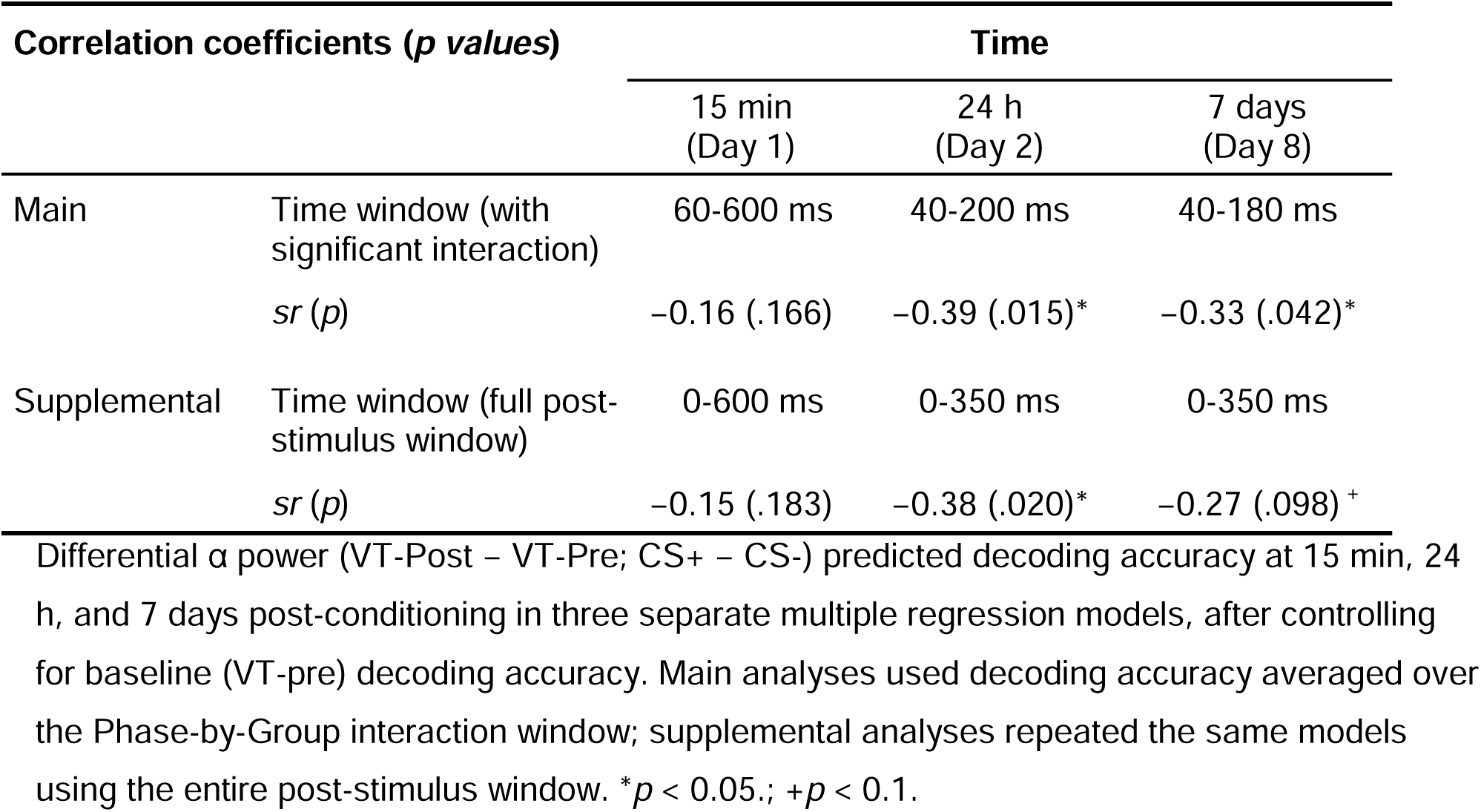
Conditioning-potentiated occipital αERD predicts decoding accuracy.

| Correlation coefficients (*p* values) |  | Time |  |  |
| --- | --- | --- | --- | --- |
|  |  | 15 min<br>(Day 1) | 24 h<br>(Day 2) | 7 days<br>(Day 8) |
| Main | Time window (with significant interaction) | 60-600 ms | 40-200 ms | 40-180 ms |
|  | *sr* (*p*) | -0.16 (.166) | -0.39 (.015)* | -0.33 (.042)* |
| Supplemental | Time window (full post-stimulus window) | 0-600 ms | 0-350 ms | 0-350 ms |
|  | *sr* (*p*) | -0.15 (.183) | -0.38 (.020)* | -0.27 (.098)<sup>+</sup> |

Differential $\alpha$ power (VT-Post – VT-Pre; CS+ – CS-) predicted decoding accuracy at 15 min, 24 h, and 7 days post-conditioning in three separate multiple regression models, after controlling for baseline (VT-pre) decoding accuracy. Main analyses used decoding accuracy averaged over the Phase-by-Group interaction window; supplemental analyses repeated the same models using the entire post-stimulus window. \* $p < 0.05$ .; + $p < 0.1$ .

## Notes

### Competing Interest Statement

The authors have declared no competing interest.

